# Identification and characterization of hPSC-derived FOXA2+ progenitor cells with ventricular cardiac differentiation potential

**DOI:** 10.1101/2021.07.18.452860

**Authors:** Damelys Calderon, Nadeera Wickramasinghe, Leili Sarrafha, Christoph Schaniel, Shuibing Chen, Mark Tomishima, Nicole C. Dubois

## Abstract

While much progress has been made in understanding early cardiac development, the precise mechanisms that specify the different cardiomyocyte subtypes remain poorly understood. Recent data from our lab have shown that transient *Foxa2* expression identifies a progenitor population with exclusive ventricular differentiation potential in the mouse heart. Here we have translated this concept to the human pluripotent stem cell (hPSC) system. Using a *FOXA2-GFP* reporter cell line we characterized expression of *FOXA2* during hPSC cardiac differentiation and found that a subset of cardiac mesoderm precursors transiently expresses *FOXA2*. Gene expression analysis of FOXA2+ and FOXA2- cardiac mesoderm revealed that both populations similarly express early cardiac specification markers such as *PDGFRA, TBX5*, and *ISL1*, while other key candidates including *TBX20* and *GATA4* are significantly upregulated in the FOXA2+ population. Isolation and subsequent differentiation of FOXA2+ and FOXA2- populations demonstrates their comparable differentiation potential to both cardiomyocytes and epicardial cells. However, cardiomyocytes derived from FOXA2+ precursors showed enhanced differentiation efficiency toward ventricular cardiomyocytes compared to cardiomyocytes derived from FOXA2- precursors. To identify new mechanisms that regulate ventricular specification, we performed small molecule screening and found that inhibition of the EGFR pathway strongly increased the cardiac mesoderm population in general, and the FOXA2+ precursors in particular. Finally, we have identified a combination of cell surface markers to specifically isolate FOXA2+ cardiac precursors. In summary, our results suggest that FOXA2+ cardiac mesoderm harbors ventricular-specific differentiation potential and isolation of these cells permits the generation of cultures enriched for ventricular cardiomyocytes. Generating such enriched cardiac populations will be relevant for regenerative medicine approaches, as well as for disease modeling from induced pluripotent stem cells.

## INTRODUCTION

The ability of pluripotent stem cells (PSC) to give rise to any cell in the human body has opened unprecedented opportunities to study human development, tissue function and homeostasis and disease^1,–3,^. Protocols to derive specific cell types of a given organ have become increasingly sophisticated, including approaches to generate the large range of the different cardiovascular cells. These include atrial and ventricular cardiomyocytes (CMs), endothelial cells, fibroblast cells, smooth muscle cells and cells of the conduction system^4,–9,^.

Recent studies have demonstrated that both atrial and sinoatrial CMs can be generated with high purity by modulating RA, FGF and WNT signaling at early stages during differentiation^4,,9,–11,^. In contrast, the protocols used to generate ventricular CMs continue to give rise to a heterogenous population of cells of which not all express canonical markers of ventricular cells^6,,8,,12,^. This suggests that either the cells of the ventricular chambers are composed of a more heterogenous mix of cells, or that current protocols are in need of further refinement to generate more defined CM populations.

Development of the heart *in vivo* is a complex process comprised of many key stages each relevant to form a properly functioning organ^13,14,^. Synthesized from different animal studies over time a concept has emerged that suggests that the early cell fate specification events are paramount for normal heart development, and that errors in generating the different progenitor populations such as the well characterized first- and second heart field cells will lead to consequences later during morphogenesis^15,–19,^.

Along these lines early fate mapping studies have suggested that chamber-specific fate is determined long before actual chamber morphogenesis, potentially as early as during gastrulation^20,21,^. Single cell sequencing and clonal lineage tracing analysis further confirmed this hypothesis, illustrating that early mesoderm populations are heterogenous and that cells that egress from the primitive streak sequentially will contribute to the future 4-chambered heart in a defined pattern, with early migrating cells contributing to FHF derivatives and later cells contributing to the SHF derivatives^17,22,23,^.

We have recently shown that transient expression of *Foxa2* during gastrulation labels a population of cells that will give rise to cardiovascular cells in the differentiated heart^24,^. Importantly, *Foxa2*-expressing cells give rise exclusively to ventricular cell types, but not atrial cells during mouse development. Here we build on these findings with the aim to identify such a ventricular-specific progenitor population during PSC differentiations *in vitro*. Using a newly generated a *FOXA2-GFP* reporter PSC line we demonstrate that FOXA2+ mesoderm are generated in vitro during cardiac differentiation, and that FOXA2+ mesoderm cell have an enhanced differentiation potential to ventricular CMs compared with FOXA2- mesoderm cells. We further uncover a role for EGFR signaling during early cardiac specification and specifically during specification of ventricular CMs. Lastly, we have identified CD148 as a cell surface marker that is specifically expressed on FOXA2+ cells. In summary, our data show that early specification events that occur in the embryo can be translated *in vitro* to generate new PSC differentiation approaches that result in highly defined cardiac cell types.

## RESULTS

### Generation of a FOXA2-GFP human PSC reporter line

We have previously identified a cardiac progenitor population in the mouse embryo that gives rise to ventricular cells, but not the atria^24,25,^. This progenitor population exists at the time of gastrulation, when it transiently expresses the transcription factor *Foxa2*. To investigate the occurrence of a complementary progenitor population during human pluripotent stem cell (hPSC) differentiation we generated a *FOXA2-GFP* hPSC reporter line using TALEN technology^26,^. A GFP construct was inserted by homologous recombination immediately following the *FOXA2* coding sequence, linked with a 2A peptide (**Fig. 1a**). This approach ensures expression of GFP under the endogenous control of the *FOXA2* regulatory elements without disrupting FOXA2 function.

**Figure 1.**
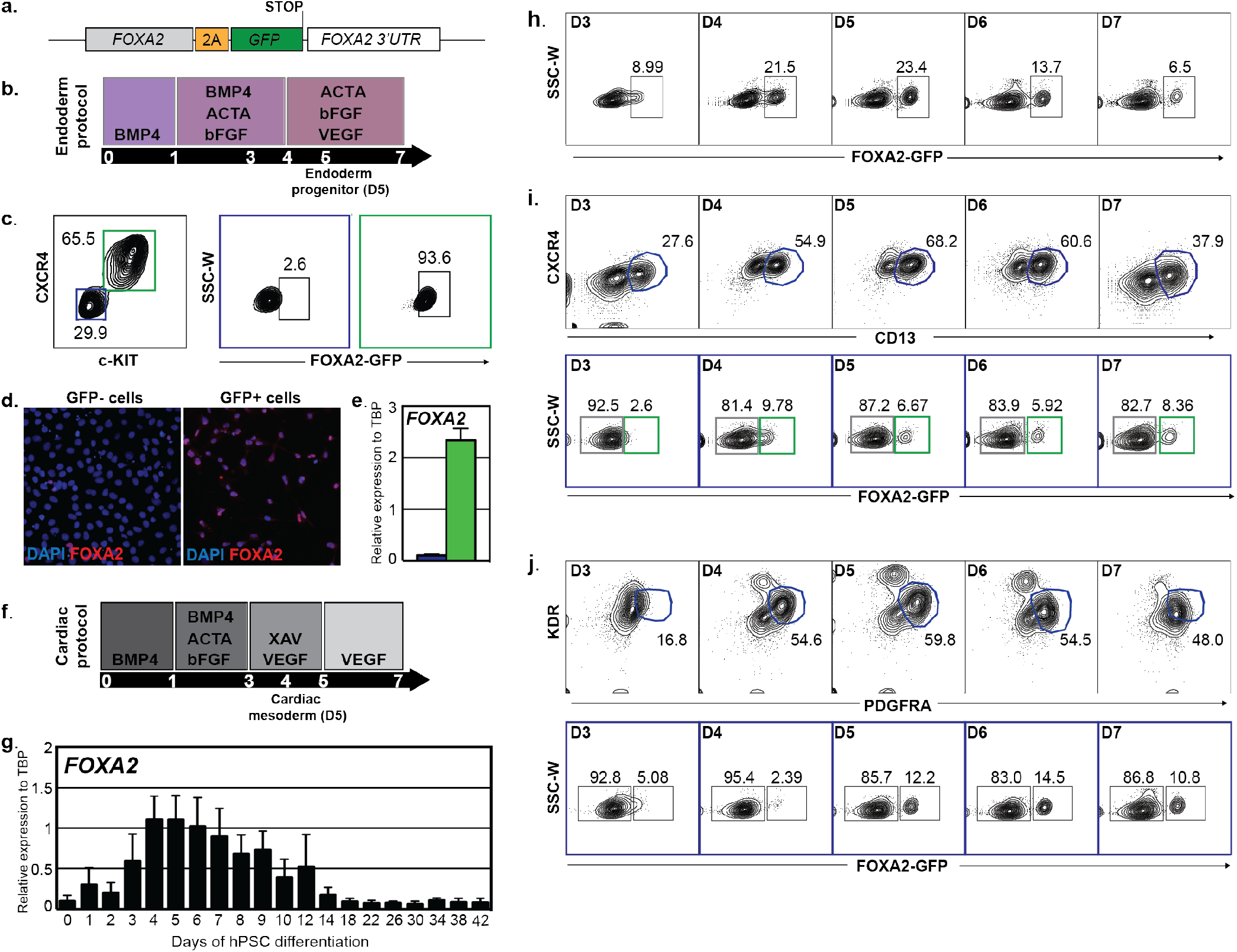
FOXA2 is expressed in a cardiac mesoderm subpopulation. **a)** Schematic of the FOXA2-GFP reporter cell line construct generated by TALEN. **b)** Schematic of the endoderm differentiation protocol performed to validate the *FOXA2-GFP* reporter cell line; **c)** Flow cytometry analysis of endoderm progenitors (CXCR4^+^c-KIT^+^) on day 5 of differentiation; **d/e)** Day 5 sorted GFP- (blue) and GFP+ (green) cells were analyzed for *FOXA2* expression by IF (d) and RT-qPCR (e); **f)** Schematic of the cardiac differentiation protocol; **g/h)** *FOXA2-GFP* expression during cardiac differentiation by gene expression (g) and flow cytometry analysis (h); **i/j)** Day 3-7 GFP expression in mesoderm population CD13+CXCR4- (i) or PDGFRA+KDR+ (j).

Correctly targeted clones were identified by PCR screening, and 5 correctly targeted clones were obtained from a total of 23 (21.7% targeting efficiency). Two clones were further transfected with Cre recombinase to excise the loxP-flanked puromycin-resistance cassette. The data shown from here on was generated using both pre- and post-Cre excision clones, and we obtained the same results with either clone.

As FOXA2 is not expressed in the heart, or in most mesoderm cells, we validated the *FOXA2-GFP* line by differentiating hPSCs to endoderm using previously described protocols (**Fig. 1b**)^27,–30,^. As expected, endoderm progenitors (marked by expression of CXCR4 and c-KIT) derived from *FOXA2-GFP* hPSCs expressed high levels of GFP, while CXCR4-c-KIT-cells did not express GFP (**Fig. 1c**). Fluorescence-activated cell sorting (FACS) of GFP+ and GFP- cells further demonstrated FOXA2 expression exclusively in GFP+ cells, confirming faithful expression of GFP in FOXA2-expressing cells (**Fig. 1d/e**).

### FOXA2 expression identifies a cardiac mesoderm subpopulation that is transcriptionally distinct compared to FOXA2-negative mesoderm

Next, we investigated FOXA2 expression during differentiation of PSCs to cardiomyocytes (CMs). hPSCs were differentiated using an embryoid body (EB) protocol previously described (**Fig. 1f**)^12,6,7,^. Quantitative PCR (qPCR) analysis over the course of differentiation illustrated a peak of transient *FOXA2* expression during the mesoderm formation stages (days 3-6)(**Fig. 1g**). Consistent with the *FOXA2* transcript expression, FOXA2-GFP+ cells were detected starting at day 3 and peaking at day 4 of differentiation using flow cytometry analyses (**Fig. 1h**).

To characterize the FOXA2-GFP+ population during cardiac differentiation, and confirm its mesodermal identity, we first performed flow cytometry analyses of lineage-specific markers at daily intervals (days 3-7) during differentiation. CD13 has been identified as a lateral plate mesoderm marker labeling MESP1+ cells, and CXCR4 was used to exclude potential endoderm cells in the culture^31,^. Our analyses revealed high CD13 expression indicative of efficient mesoderm formation, as well as a distinct FOXA2-GFP+ population within the CD13+ mesoderm (**Fig. 1i**). KDR and PDGFRA specifically mark cardiac mesoderm^7,9,10^. As expected, high levels of KDR+PDGFRA+ cells were detected during the mesoderm forming stages of differentiation, and FOXA2-GFP+ cardiac mesoderm cells were present as early as Day 3 of differentiation (**Fig. 1j**). These data suggest that FOXA2+ mesoderm cells are generated during human cardiac development *in vitro*, similar to what has been observed during *in vivo* and *in vitro* mouse development^24,^.

To further confirm the mesoderm identity of the FOXA2-GFP+ population generated during cardiac development, CD13+CXCR4-GFP+ and CD13+CXCR4-GFP- cells were isolated by fluorescence-activated cell sorting (FACS) and subjected to immunofluorescence and gene expression analyses at day 4 of differentiation (**Fig. 2a-c**). Sorted CD13+CXCR4-GFP+ and CD13+CXCR4-GFP- populations showed high expression of MESP1 (90% and 78% respectively), but only CD13+CXCR4-GFP+ cells co-expressed FOXA2 (**Fig. 2b/c**). hPSC-derived endoderm progenitors (CXCR4+c-KIT+) were used as a positive control for FOXA2 expression. As expected we detected high expression of FOXA2, but not MESP1 in endoderm cells (**Fig. 2b/c**).

**Figure 2.**
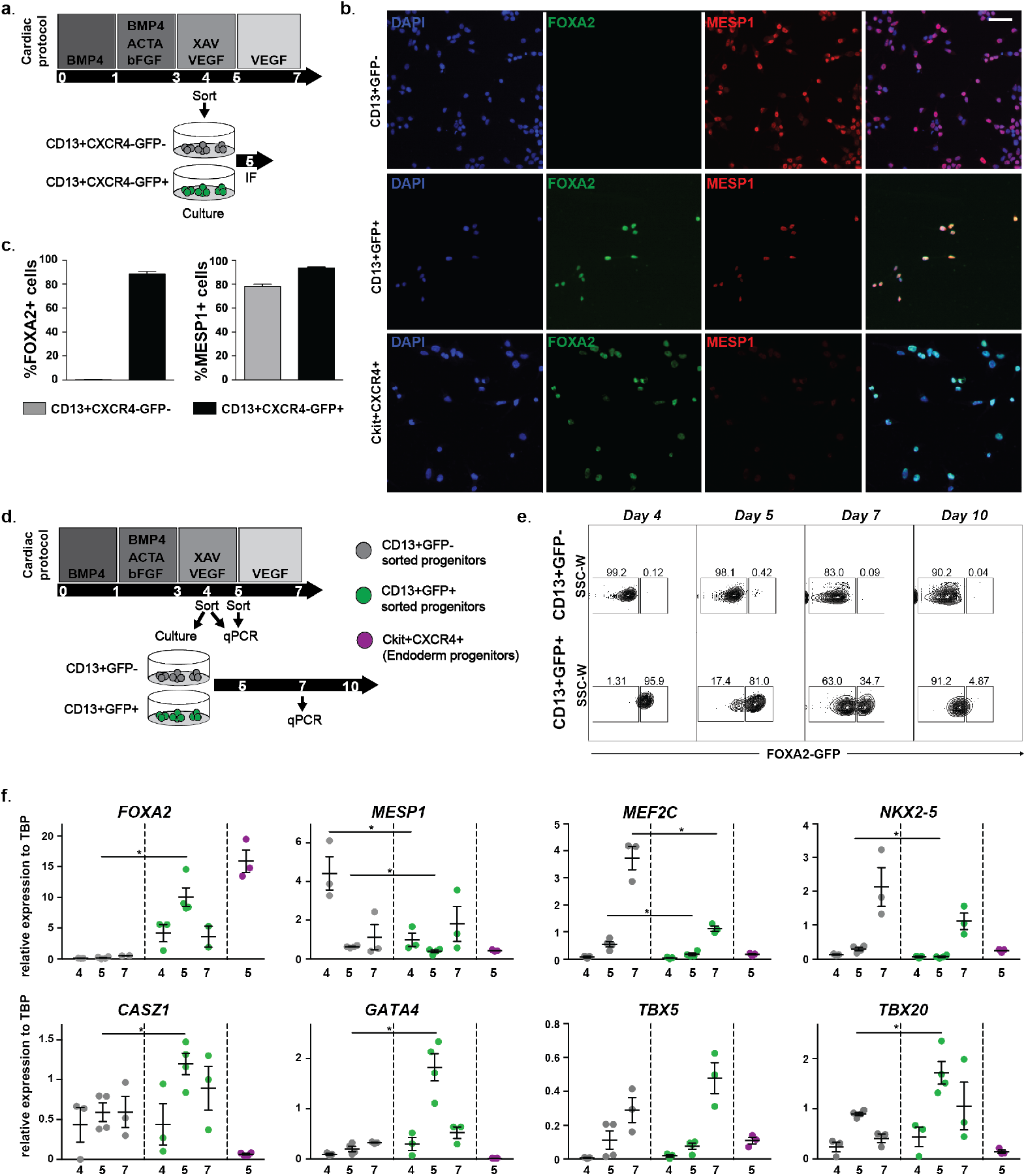
FOXA2+ mesoderm shows differential gene expression compared to FOXA2- cardiac mesoderm. **a/d)** Schematic of the cardiac differentiation protocol and sorting strategies. **b/c)** GFP+ and GFP- mesoderm and endoderm progenitors were sorted on D4 (mesoderm) or D5 (endoderm) and analyzed for *MESP1* and *FOXA2* expression by IF (b) with quantification in (c); **e)** GFP+ and GFP- mesoderm cells were sorted at day 4 of differentiation and cultured separately, and GFP expression was analyzed by flow cytometry on day 5, 7 and 10 in both populations; **f)** RT-qPCR analysis of sorted GFP- (grey) and GFP+ (green) mesoderm populations on day 4, 5 and 7 and endoderm progenitors (violet) on day 5. Graphs represent the relative mean expression value and SEM normalized to TBP from three independent differentiations. Statistical analysis was performed only between mesoderm populations on the same day.

In the mouse embryo, Foxa2 expression during cardiac development is transient and not detected anymore after embryonic day 8.50 (E8.5)^24,^. To monitor FOXA2 expression during human cardiac development *in vitro*, CD13+CXCR4-GFP+ and CD13+CXCR4-GFP- cells were sorted, cultured, and GFP+ cells were monitored (**Fig. 2d**). While CD13+CXCR4-GFP- cells never up-regulated GFP, 90% of the CD13+CXCR4-GFP+ cells had lost GFP by day 10 of differentiation, demonstrating that FOXA2 expression in cardiac mesoderm is transient, as has been observed during mouse heart development (**Fig. 2e**). This data further indicates that FOXA2 expression is restricted to the early mesoderm stages and is not observed at later time points during differentiation.

Gene expression analyses at different times during early differentiation showed that both CD13+CXCR4+GFP- and CD13+CXCR4-GFP+ cells expressed comparable levels of key genes relevant for the formation of the cardiac lineage (*MESP1, PDGFRA NKX2-5, MEF2C* and *TBX5*), and that the expression levels were higher compared to endoderm progenitors (**Fig. 2f**). Interestingly, we detected differences in gene expression levels of some of the early cardiac genes (*TBX20* and *GATA4*) between CD13+CXCR4-GFP- and CD13+CXCR4-GFP+ cells, potentially indicating that the FOXA2- and FOXA2+ cardiac lineages are distinct as early as at the mesoderm stage. In support of this, *CASZ1* and *FOG2*, two genes previously found to be upregulated in the Foxa2+ mouse mesoderm were also upregulated in FOXA2+ human mesoderm (**Fig. 2f**, **Supplementary Fig. 1**)

### FOXA2+ mesoderm preferentially differentiates to ventricular cardiomyocytes

The central objective of this study was to identify the ventricular-specific Foxa2+ progenitor population found *in vivo* during mouse development in *in vitro* human differentiations, in order to establish robust protocols to generate pure ventricular cell types. To characterize their cardiac differentiation potential, CD13+CXCR4-GFP- or CD13+CXCR4-GFP+ cells were sorted (day 4) and cultured as aggregates in suspension for two additional months (**Fig. 3a**). At day 60, the efficiency of CM differentiation was quantified by flow cytometry analyses for SIRPA (CMs) and CD90 (fibroblast-like cells)^7,^. We found that cardiomyocytes were generated from both GFP- and GFP+ mesoderm populations with equal frequency, further confirming the mesodermal identify of CD13+CXCR4-FOXA2+ cells (**Fig. 3b/c**). Un-sorted cardiomyocytes were used as a reference.

**Figure 3.**
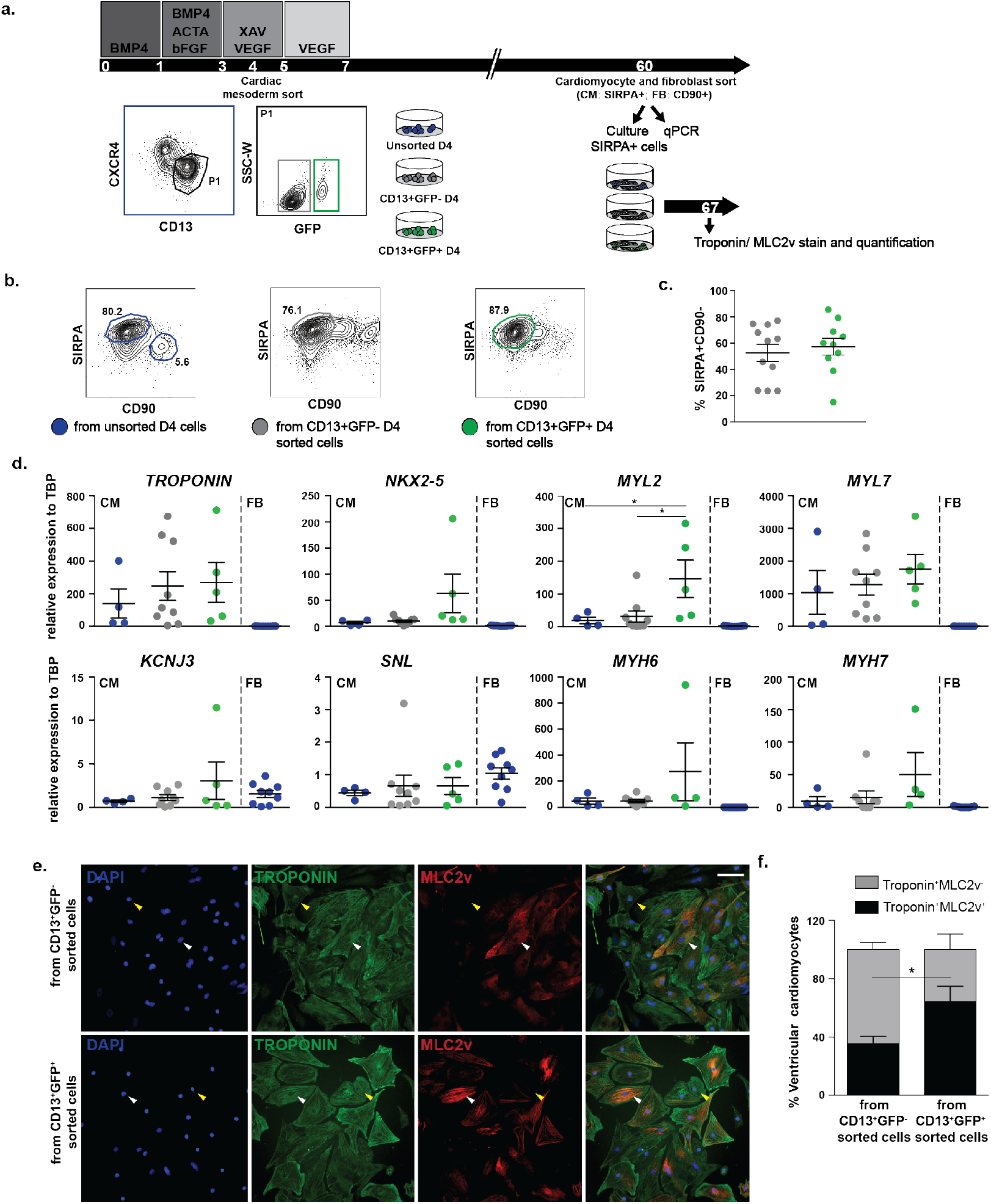
FOXA2+ mesoderm preferentially differentiates to ventricular cardiomyocytes. **a)** Schematic of the cardiac differentiation protocol and sorting strategy. GFP- (grey) and GFP+ (green) mesoderm cells were sorted at day 4 of differentiation and cultured separately. On day 60 of differentiation SIRPA-CD90+ (FB) and SIRPA+CD90- (CM) were sorted from each fraction or from whole differentiations (bleu); **b)** Representative SIRPA/CD90 profiles and their quantifications (n=11); **c)** Gene expression analysis of sorted populations at day 60; **d)** MLC2v and Troponin T IF staining of sorted CM populations at day 60, and the quantitative analysis of MLC2v+/Troponin+ and of MLC2v-/Troponin+ phenotypes in culture at day 60. The data shown represent the mean ± the SEM from two independent differentiations.

To determine whether FOXA2-GFP- and Foxa2-GFP+ mesoderm cells give rise to different CMs, we performed gene expression analyses on SIRPA+ cells two month after differentiation. This analyses revealed comparable expression of sarcomere-related transcripts *TNNT2, MYH6, MYH7* and of the pan-cardiac transcription factor *NKX2-5* in FOXA2-GFP- and FOXA2-GFP+-derived CMs. Atrial markers *KCNJ3* and *SNL* showed equal expression in both populations. By contrast, the ventricular marker *MYL2* (encoding MLC2v) was expressed 5-fold higher in the CD13+CXCR4-FOXA2:GFP+ compared to the CD13+CXCR4-FOXA2:GFP- derived or unsorted CMs (**Fig. 3d**). The enrichment of ventricular CMs in FOXA2-derived cultures was confirmed by IF analyses, demonstrating that significantly greater percentage of MLC2v+TroponinT+ cells in FOXA2-GFP+ derived cultures compared to FOXA2-GFP- derived cells (**Fig 3e/f**). These data suggest that indeed the *in vivo* Foxa2+ ventricular progenitor population can be identified in culture, and that similar to the mouse embryo, it gives rise to preferentially ventricular cardiomyocytes during differentiation.

### EGF pathway inhibition during early mesoderm differentiation increases the generation of FOXA2+ cardiac mesoderm

In contrast to the mouse embryo, the FOXA2+ cardiac mesoderm population generated during human *in vitro* differentiation with the current protocol is comparatively small (**Fig. 1h-j**). To identify signaling pathways that regulate, and potentially increase the generation of FOXA2+ cardiac mesoderm we performed a small molecule screen during the mesoderm induction stage. We assembled a screen composed of 208 small molecules targeting the major pathways relevant for early development and cell fate specification, including but not limited to WNT, SHH, FGF, PI3K. The compounds (final concentration of 10uM) were added individually to cells from day 1-3 of differentiation, and the cells were evaluated on day 4 of differentiation for the generation of FOXA2+ mesoderm (**Fig. 4a/b and Supplementary Fig. 2**). From this screen we identified 17 candidates that resulted in increased percentages of CD13+FOXA2-GFP+ cells. Cells treated with these compounds were then further differentiated using a cardiac differentiation protocol and analyzed by flow cytometry for SIRPA expression to confirm their ability to still efficiently generate CMs (**Fig. 4c**). Next we assessed the effects of these compounds on the generation of ventricular CMs by IF analysis of the ventricular-specific marker MLC2v (**Fig. 4d**). From this combined analysis we defined 4 categories with distinct phenotypes: *category I*: compounds that increase FOXA2+ cardiac mesoderm and ventricular CM generation; *category II*: compounds that do not change FOXA2+ mesoderm and increase ventricular CM generation; *category III*: compounds that increase FOXA2+ cardiac mesoderm and do not increase ventricular CM generation and *category IV*: compounds that that decrease FOXA2+ cardiac mesoderm and decrease ventricular CM generation (**Fig. 4b/d**). IF analysis for cardiac Troponin T and MLC2v is shown for a select example in each category (**Fig. 4d**).

**Figure 4.**
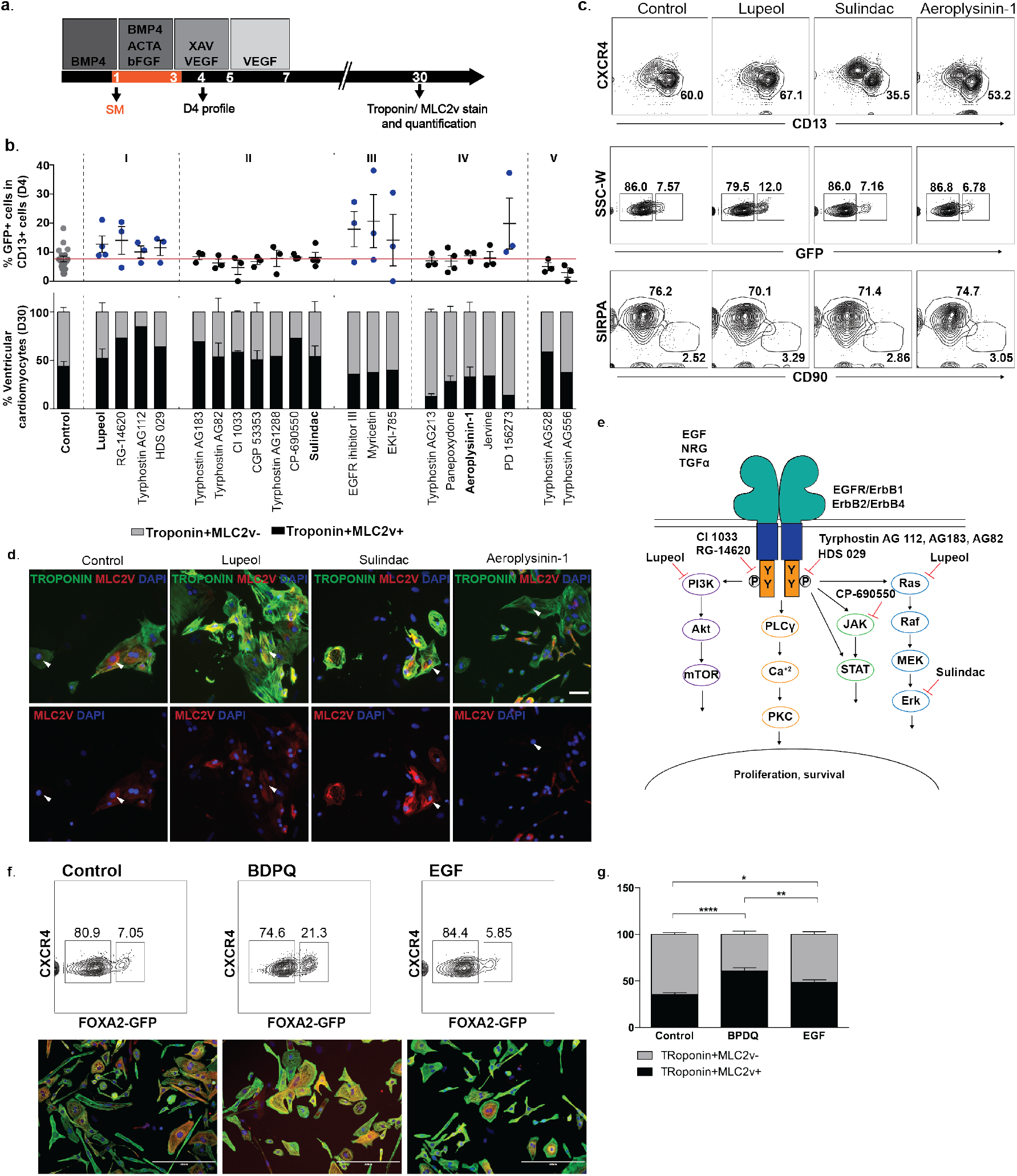
EGF pathway inhibition at the mesoderm induction stage increases generation of FOXA2+ cardiac mesoderm. **a)** Schematic of the cardiac differentiation protocol and the small molecule screen strategy. A total of 208 SM were individually added at day 1 of differentiation; **b)** Quantification of percentages of GFP expressing cells on day 4 mesoderm population (CD13+) and quantification of Troponin and MLC2v expression on day 30. Group classification (I to V) is determined by these two parameters; **c)** Flow cytometry analysis of D4 mesoderm populations (CD13, CXCR4 and GFP) and cardiomyocytes (SIRPA,CD90) treated with small molecules on D1 (Lupeol, Sulindac and Aeroplysinin-1) or treated with DMSO only (control); **d)** IF analysis on day 30 cardiac cultures from SM treated cultures and control. Quantitative analysis of MLC2v+/Troponin+ and of MLC2v- /Troponin+ phenotypes are shown in (b) and represents the mean ± the SEM from two independent differentiations; **e)** Schematic representation of the epidermal growth factor receptor (EGFR) and ERBB proteins and their downstream pathways. Small molecules that increased the generation of ventricular cardiomyocytes (group I and II) inhibited the EGFR autophosphorilation process or their downstream pathways. **f)** Flow cytometry analysis of FOXA2+ in mesoderm populations at day 4 of differentiation after EGF actovation/inhibition (upper panles) and IF analysis at day 60 of differentiation for MLC2v and Troponin T; **g)** Quantification from (f).

Interestingly, all the small molecules that resulted in increased ventricular CM generation (categories I and II) negatively target the EGFR pathway, either upstream by inhibiting autophospohorylation of the EGFRs (RG-14620, Tyrphostin AG112, HDS029, Tyrphostin AG183, Tyrphostin AG82, CI 1033), or downstream by targeting the different branches of EGFR signaling (PI3K, JAK and ERK pathways)(**Fig. 4e**). To follow up on this we evaluated the expression of individual EGFRs over the course of human *in vitro* cardiac differentiation. RT-qPCR analysis illustrates that EGFR1 and EGFR2 are expressed a low levels during hPSC differentiation, however, EGFR3 and EGFR4 are highly expressed during the mesoderm formation stage, indicating that inhibition of their signaling activity at this stage may have distinct effects (**Supplementary Fig. 3**). To determine whether modulation of the EGF pathway can impact atrial-ventricular fate from hPSCs we treated cardiac progenitor cells with either recombinant EGF (50-100ng/ml) or with BDPQ (EGFR inhibitor) for 48hrs (days 1-3) and analyzed cultures for FOXA2-GFP expression in mesoderm at day 4 of differentiation. No decrease of FOXA2 expression was observed in the mesoderm populations in the presence of EGF, and no decrease in ventricular cells was observed in differentiated CMs, but inhibition of EGFR signaling with BDPQ increased both the FOXA2+ mesoderm population and differentiated MLC2v+ CMs (**Fig. 4f/g**).

In summary, using the FOXA2+ cardiac mesoderm population as a read-out for ventricular CM potential we have identified the EGFR pathway as a modulator for the early specification of the ventricular lineage in human *in vitro* CM development.

### Inhibition of retinoic acid signaling does not impact the formation of FOXA2+ mesoderm during hPSC differentiation

Animal models and more recently work in hPSC differentiations have determined that retinoic acid (RA) signaling patterns cardiac progenitor cells along the anterior-posterior axis of the embryo, and that RA plays a role in the induction of the atrial CM fate^4,,32,–37,^. These data further illustrate that cardiac specification is a process that occurs early during development, at the mesoderm and progenitor stages, and that RA signaling activation is sufficient to induce an atrial-like phenotype in hPSC-derived CMs. In order to investigate whether and when the RA pathway is active during hPSC differentiation, we first monitored expression of Retinaldehyde dehydrogenase 2 (*RALDH2*), the enzyme that oxidizes retinal to retinoic acid. *RALDH2* was found to be expressed in a bi-phasic pattern, with one wave occurring early during differentiation (correlating with mesoderm formation) and a second wave occurring at the stage after differentiation to cardiomyocytes is completed (**Fig. 5a**). Furthermore, *Cyp26a1* a key enzyme in the pathway that facilitates degradation of RA expression was also assessed, as it has been shown that Cyp26-deficient zebrafish embryos cannot maintain the integrity of the nascent heart tube because of the ventricular cells losing their polarity. We find that *Cyp26a1* is strongly upregulated on day 3 of differentiation (early mesoderm) and very rapidly downregulated after that and remains low for the duration of cardiac differentiation thereafter (**Fig 5a**). These gene expression patterns corelate closely with the notion that RA signaling plays an important role during early cardiac specification. To test whether hPSCs can be differentiated toward a ventricular phenotype, as previously described, EBs were treated with BMS-189453 (a pan-retinoic acid receptor antagonist) from day 5 to 7 at a concentration of 10 μM. At day 45, EBs showed an increased percentage of cells expressing *MLC2v* in the RAi treated cultures compared to the controls (**Supplementary Fig. 4**). We next inhibited RA signaling during mesoderm formation (days 3-5), which led to a small increase in FOXA2+ cardiac mesoderm early during differentiation (day 4)(**Fig. 5b/c**). We next sought to determine the effect of RA inhibition in FOXA2- and FOXA2+ mesoderm separately. FOXA2- and FOXA2+ mesoderm was sorted at day 4 of differentiation, cultured in the presence of BMS-189453 for 48hrs followed by additional culture for 54 days (**Fig. 5d**). Flow cytometry analysis for SIRPA and CD90 revealed that inhibition of RA signaling in FOXA2+ mesoderm resulted in increased differentiation efficiency toward cardiomyocytes compared to FOXA2- cells (**Fig. 5e-f**). Moreover gene expression analysis of GFP+ and GFP- derived cardiomyocytes from both control (no BMS-189453, blue dots) and BMS-189453 treated populations (GFP- mesoderm derived cardiomyocytes, grey; GFP+ mesoderm derived cardiomyocytes, green) confirms that BMS-189453 treatment affects the generation of cardiomyocytes (**Fig 5g**).

**Figure 5.**
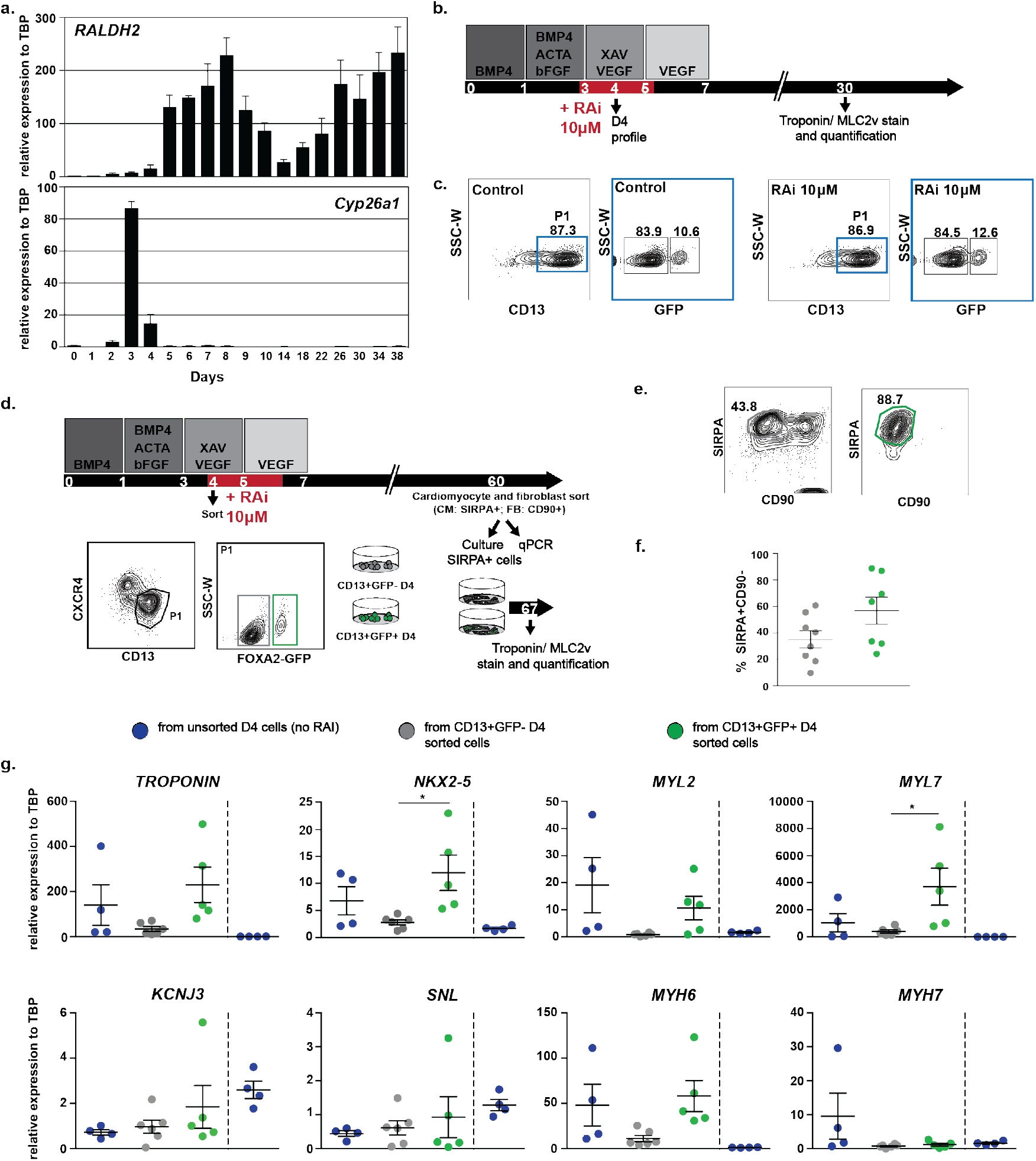
Inhibition of retinoic acid during cardiac ventricular specification. **a)** RT-qPCR expression of *RALDH2* and *Cyp26a1* during cardiac differentiation; **b)** Schematic of the cardiac differentiation protocol and the retinoic acid inhibition (RAi) strategy; **c)** Flow cytometry analysis of D4 mesoderm populations (CD13, CXCR4 and GFP) with RAi or DMSO only (control); **d)** Schematic of the cardiac differentiation protocol and sorting strategy. GFP- (grey) and GFP+ (green) mesoderm cells were sorted at day 4 of differentiation and cultured separately in RAi until day 6. On day 60 of differentiation SIRPA-CD90+ (FB) and SIRPA+CD90- (CM) were sorted from each fraction; **e/f)** Representative SIRPA/CD90 profiles and their CM percentages (n=7); **g)** Gene expression analysis of sorted populations at day 60. The data shown represent the mean ± the SEM from 5 independent differentiations.

### CD148 is a specific cell-surface marker for isolating FOXA2 expressing cells

In order to find a broadly applicable approach to isolate FOXA2+ mesoderm without having to rely on a reporter line we performed flow cytometry screening with a library of cell surface antibodies. 369 known surface antibodies were used on mesoderm derived from the *FOXA2-GFP* PSC line. The screen focused on identifying antibodies that recognized antigens present on the CD13+CXCR4-GFP+ population. CD148, a receptor tyrosine phosphatase, was the only one that displayed a FOXA2-specific expression pattern (**Fig 6a**). These results were confirmed by using cells derived from H7 PSCs (**Fig 6a**). CD148 expression was also observed on D5 endoderm progenitors using the reporter FOXA2-GFP cell line and the wild type H7 cell line, confirming its overall specificity to FOXA2- expressing cells (**Fig 6b**). To confirm that CD148 can be utilized to sort FOXA2+ ventricular progenitor cells from H7-derived cultures, hPSCs were differentiated to cardiac mesoderm and sorted on day 4 for CD13+CXCR4-CD148+ or CD13+CXCR4-CD148- cells (**Fig. 6c**). Gene expression analysis confirmed *FOXA2* expression in CD148+ cells from both mesoderm and endoderm differentiations, while CD148- mesoderm was negative for *FOXA2* (**Fig. 6d**). Similarly, IF analysis 24hrs after sorting confirmed FOXA2 expression in CD13+CXCR4-CD148+ cells, which also co-expressed the mesoderm marker MESP1 (**Fig 6e**). Collectively our data show that CD148 is a cell surface marker specifically expressed on FOXA2 expressing cell types, and that sorting for CD148 during cardiac differentiations enable isolation of ventricular progenitor cells similar to using the FOXA2-GFP reporter line.

**Figure 6.**
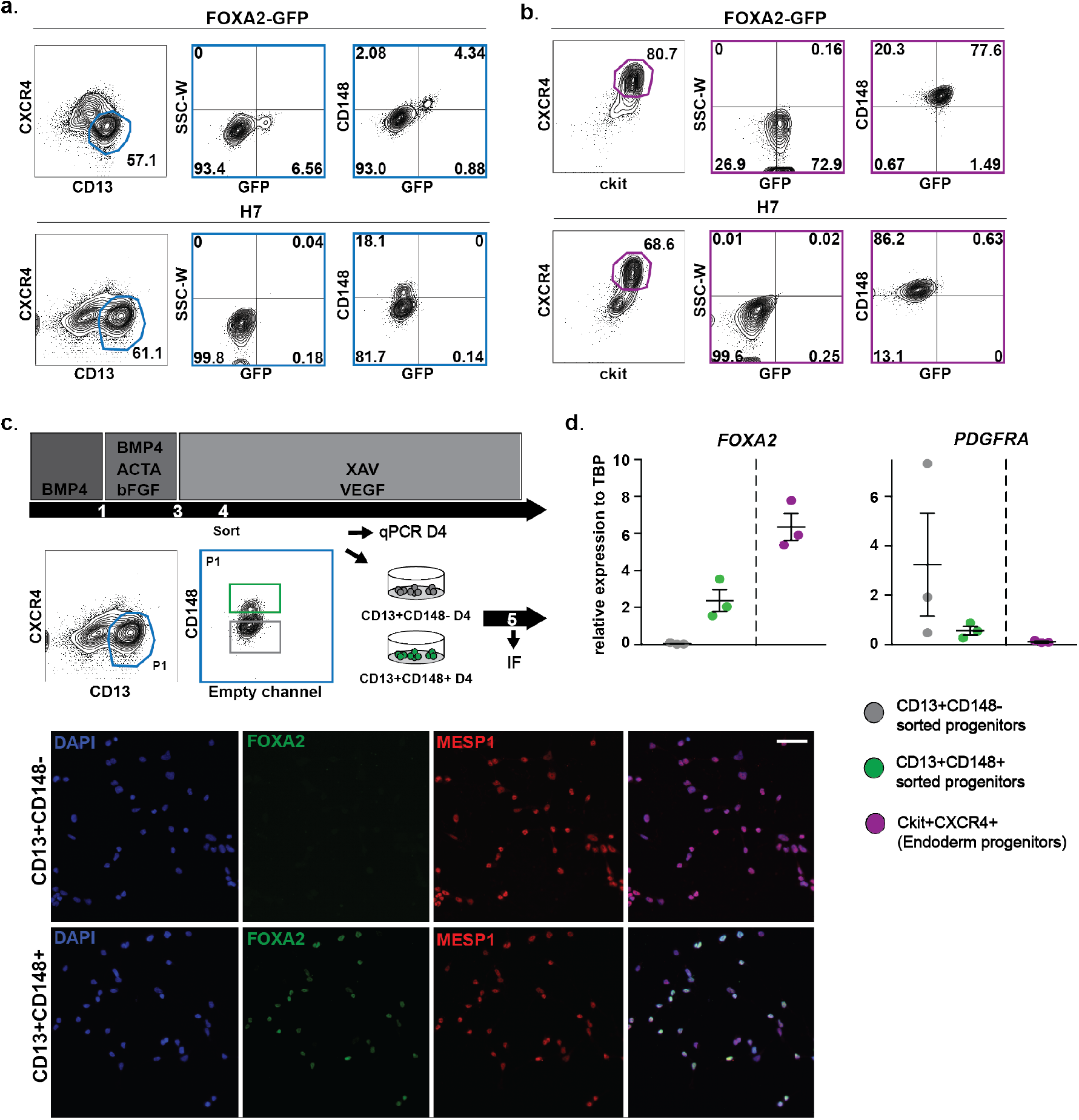
CD148 is specifically expressed in FOXA2+ cardiac mesoderm and endoderm. **a/b)** Mesoderm differentiations (a) and endoderm differentiations (b) using the *FOXA2-GFP* reporter cell line (top) or a control cell line (H7, bottom) showes CD148 expression in GFP+ mesoderm and endoderm cells. Similar profiles were observed when using control cell lines; **c/d)** Schematic of the cardiac differentiation protocol and sorting strategies (c). CD148+ (green) and CD148- (grey) mesoderm progenitors were sorted on D4 and analyzed for FOXA2 and PDGFRA expression by qPCR (d) and their levels of expression compared to sorted endoderm progenitors (violet); **e)** D4 sorted mesoderm populations were also cultured separately one day and analyzed by IF for FOXA2 and MESP1 expression. The data shown is representative of three independent experiments.

## DISCUSSION

Work in both animal models and PSC differentiations suggests that cardiac specification begins early during development. Early fate mapping studies have identified the prospective cardiac lineage as early as during formation of the primitive streak^20,^. Single cell sequencing and clonal lineage tracing further revealed that early cardiac progenitor cells are heterogenous and harbor their future fate in the differentiated heart already at stages prior to chamber morphogenesis^23,22,17,16,^. Along similar lines we have previously shown in the mouse that Foxa2 is transiently express in cardiovascular progenitors during gastrulation, and that Foxa2-expressing cells will give rise exclusively to cells of the ventricular chambers^24,^. Here, we demonstrate that transient FOXA2 expression can also be observed in the *in vitro* human pluripotent stem cell system, and that such FOXA2+ mesoderm shows preferential differentiation potential to the ventricular lineage.

In the mouse Foxa2 is expressed prior to the formation of the well characterized progenitor populations such as cardiac mesoderm or cells of the first and second heart fields. PSC differentiation cultures derived using standard protocols are well known to be highly heterogenous, including ventricular, atrial and conduction system cell types^8,12,^. Protocols to specifically generate atrial and sinoatrial nodal cells have recently described, however, protocols to specifically generate pure populations of ventricular cardiomyocytes are still lacking^4,,9,,10,,35,^. Our data from the mouse suggests that ventricular-specific progenitor cells do exist during development, and that FOXA2 might be a marker that can be utilized to identify such progenitor cells. In order to characterize such a potentially corresponding population *in vitro* we examined hPSC differentiation cultures for co-expression of one of the earliest markers for mesoderm formation, CD13, in combination with FOXA2 using a newly generated *FOXA2-GFP* reporter line^31,^. CXCR4 was included as an additional marker to exclude primitive endoderm which is known to also express FOXA2. Using this strategy we found that FOXA2 is indeed expressed in a sub-population of mesoderm cells that co-express CD13 and are negative for CXCR4, suggesting that the population previously discovered during mouse development can be identified during hPSC differentiations *in vitro*. Moreover, we found that FOXA2+ mesodermal progenitors showed enhanced differentiation potential towards the ventricular lineage, as evidenced by gene expression and IF analysis for canonical ventricular markers such as MLC2v.

One of the few pathways that is known to determine atrial versus ventricular fate is retinoic acid (RA) signaling. RA is expressed as a gradient during cardiac progenitor formation, and higher expression of RA favors the formation of atrial cardiomyocytes both *in vivo* and *in vitro*. In the mouse, retinaldehyde dehydrogenase 2 (*Raldh2*) expression is observed exclusively in sino-atrial tissues while ventricular tissue remained devoid of RA metabolism, again suggesting that RA is necessary for atrial development, but not for the development of ventricular cells. We had hypothesized that inhibition of RA signaling will further enhance ventricular CM differentiation, and that this effect would be more pronounced in FOXA2+ mesoderm. RA inhibition did indeed lead to an increase in ventricular CMs, and that effect seemed more pronounced in FOXA2+ derived CMs compared to FOXA2- derived CMs, suggesting that a combination of cell isolation strategies and additional signaling pathway modulations may be required to generate pure ventricular CM populations.

Having a read out for differential mesoderm populations at hand with the FOCA2-GFP reporter line we sought to use this to identify mechanisms that are at play during the generation of FOXA2+ mesoderm. As such we identified by means of a small molecule screening that inhibition of the EGFR pathway and its downstream targets (like the PI3K, JAK and ERK pathways) increase the generation of FOXA2+ mesoderm cells *in vitro*. This corroborates data from the mouse that has shown that Erbb receptors are important during cardiac morphogenesis. Erbb2 and 4 are both expressed in ventricular CMs and their disruption results in a lack of ventricular trabeculation. However their role during early cardiac cell fate specification has not been broadly studied. Our gene expression studies confirm the importance of EGFR inhibition in the generation of FOXA2+ mesoderm mechanistically, as EGFR inhibition led to upregulation of *FOG2* in FOXA2+ mesoderm compared to the FOXA2- mesoderm population, which in turn acts as a negative modulator of the PI3K-Akt pathway via direct binding to p85alpha. Multiple important roles for *FOG2* have been described during myocardial development by generating conditional knockouts in CMs^38,39,^. Lastly, erbB1 signaling was shown to be able to regulate the retinoic acid pathway, providing a possible sequence of events during early cardiac specification^40,^.

Finally, to broaden the approach to any cell line of interest our cell surface screening experiments identified CD148 is a specific cell-surface marker for isolating FOXA2 expressing cells in cardiac mesoderm and endoderm. CD148 is a member of the receptor tyrosine phosphatase family (*PTPRJ*), and is known to regulate maturation of primary cellcell contacts, endothelial proliferation and endothelium-pericyte interactions. An important role has also been reported during heart development. CD148ΔCyGFP homozygous embryos show decreased thickness of the myocardial wall, coupled with increased space between the endocardium and myocardium, disorganized trabeculation, discontinuous endocardia and lacked mesenchymal cushion formation adjacent to the atrioventricular canal^41,^. Interestingly, a binding site for FOXA2 transcription factor has been identified upstream the transcription start site of the PTPRJ gene, which indicates that *FOXA2* might be a direct regulator of PTPRJ.

In conclusion, we shoed that a FOXA2+ mesoderm population found in mouse embryos is present in hPSC cardiac differentiations and that it shows a higher ventricular cardiac potential compared to FOXA2- mesoderm. Enrichment of this population can be achieved by inhibiting EGFR signaling early during differentiation and FOXA2+ cells can be isolated from any PSC line combining antibodies against the CD13 and CD148 cell surface proteins.

## MATERIALS AND METHODS

### Construction of TALENs and donor vector to generate a FOXA2 reporter cell line

The software used to design TALENs in this study (Zifit.partners.org) uses assembly protocols described by the Joung lab (*Sander et al., Nat Biotechnol. 2012, Reyon et al. 2012 Nat Biotechnol, Reyon et al. 2012 Current Protocols in Molecular Biology*) to define the optimal sequences for potential TALE transcriptional activator/repressor target sites. TALENs were directed against the 3’ end of the FOXA2 gene. To generate TALENs, we cloned the DNA encoding the TALE repeats and flanking regions into JDS74 (plasmid #32288) and JDS71 (plasmid #32287) vectors (Addgene). The vector OCT4-2A-eGFP-PKG-Puro (plasmid #31938, Addgene) was used as template to generate the donor vector. The left and right *FOXA2* homology arms, (885 base pairs) and (300 base pairs) respectively, were amplified from human genomic DNA and clone into donor vector by means of *SbfI* and *NheI*, or *NotI* and *AscI* digestion and ligation (respectively). Correct insertion was verified by sequencing prior to genome targeting.

### Transfection of TALENs and donor vector into H7 human embryonic stem cells

H7 embryonic stem cells were transfected with the indicated TALE constructs and donor vector (at ratio 0.5μg left Talen: 0.5μg right Talen: 1.7μg donor vector per well) in Lipofectamine 2000 (Invitrogen) according to the manufacturer’s recommendations. At 96 hours after transfection, medium with selective antibiotic (puromycin 0.5 μg/ml) was added to cells for 5 days. Individual colonies were then transferred to smaller wells for initial expansion on matrigel and mTeSR medium. DNA from clones was isolated using purelink genomic DNA mini kit (K1820-01, Invitrogen) and PCR to test correct insertion of the GFP was performed. 21.7% of the clones showed correct insertion.

Removal of PGK-Puro cassette by transient Cre-recombinase expression (Vector pCAG-Cre #13776, Addgene) was performed on clones with correct GFP insertion, following similar transfection protocol previously detailed. At 48 h post-transfection single GFP expressing cells were sorted and plated onto matrigel coated 6 well plate (100.000 cells/well) containing mTeSR medium. Single cell clones were allowed to proliferate and analyzed by PCR for correct cutting. 80% of cells showed correct cutting.

### Human FOXA2-GFP ESC endoderm differentiation

FOXA2-GFP reporter cell line was differentiated into the endoderm lineage as previously described (Goldman et al., 2013). Briefly, cells were dissociated using collagenase B (Roche) and cultured in low cluster plates to allow EB formation in serum-free differentiation SFD media supplemented with BMP4 (3 ng/ml, R&D Systems). At day 1 the medium was changed to SFD media supplemented with Activin A (100 ng/ml, R&D Systems), basic FGF (bFGF, 2.5 ng/ml, R&D Systems), and BMP4 (0.5 ng/ml, R&D Systems). At day 4, the medium was changed to the SFD media supplemented with Activin A (100 ng/ml), bFGF (2.5 ng/ml), and vascular endothelial growth factor (VEGF, 10 ng/ml, R&D Systems). On day 5 EBs were dissociated, and sorted CXCR4+ cKit+ cells were used for qPCR or plated on matrigel-coated coverslips in D4 media and used for IF.

### Human FOXA2-GFP ESC cardiac differentiation

Embryoid bodies (EBs) were generated from ESC lines using collagenase B (Roche) and cultured in base media (RPMI 1640 (Invitrogen) medium containing 0.5X B27 supplement (Life technologies), 2mM glutamine (Gibco-BRL), 1 mM ascorbic acid (Sigma), and 4×10-4 M monothioglycerol (Sigma)) with BMP4 (1 ng/ml, R&Dsystem) and Thiazovivin (2 microM, Millipore). On day 1, EBs were harvested and resuspended in induction medium (base media with basic fibroblast growth factor (bFGF; 2.5 ng/ml, R&Dsystem), activin A (20 ng/ml, R&Dsystem) and BMP4 (20 ng/ml)). On day 3, the EBs were harvested and resuspended in base media with vascular endothelial growth factor (VEGF; 10 ng/ml, R&Dsystem) and XAV939 (10microM, Stemolecule). On day 8 and thereafter, EBs were cultured in base media till day 60. EBs from day 3 to day 7 were analyze by FACS and qPCR. On day 60, SIRPA+CD90- and SIRPA-CD90+ cells were sorted and used for qPCR or plated on matrigel-coated coverslips in base media with Thiazovivin (2 microM) and culture for 7 days before IF analysis.

### FOXA2 cardiac progenitor differentiations

On day 4 of cardiac differentiation, EBs were dissociated using TrypLE (Invitrogen). CXCR4-CD13+GFP+ and CXCR4-CD13+GFP- cells were sorted and used for mRNA extraction, IF analysis or cultured in pluronic-treated 96 well plates at 1 × 10^5^ cells per well in base media with vascular endothelial growth factor (VEGF; 10 ng/ml), XAV939 (10microM) and Thiazovivin (2 microM). On day 5, half of the media was changed to base media with vascular endothelial growth factor (VEGF; 10 ng/ml), and cultures were maintained in this media until day 8 of differentiation, when EBs were culture in base media only, with media changes every 3 days till day 60, where cells were analyzed as previously described.

### Small molecule screen

On day 1 of cardiac differentiation FOXA2-GFP EBs were plated into pluronic-treated 96 well plates at 1 × 10^5^ cells per well in day 1 cardiac differentiation media and a single compound from a library containing 208 small molecules (Suplementary table 1), at a 1:1000 dilution. Differentiation was then carried out as previously described. Cardiac mesoderm profile (CD13 and CXCR4 stain) was performed on day 4 of differentiation. Two criteria were used to define the primary hits compounds for further analysis. On day 30, cells were dissociated and analyzed for SIRPA and CD90 expression and plated on matrigel-coated 96 well plate in base media with Thiazovivin (2 microM) and culture for 7 days before IF analysis.

### Flow cytometry and cell sorting

#### Dissociation procedure for day 3 to day 12 EBs

EBs generated from hPSC differentiation experiments were dissociated with 0.25% trypsin/EDTA.

#### Dissociation procedure for day 13 and older EBs

EBs were incubated in collagenase type II (1 mg/ml; Worthington) in Hanks solution (NaCl, 136 Mm; NaHCO3, 4.16 mM; NaPO4, 0.34 mM; KCl, 5.36 mM; KH2PO4, 0.44 mM; dextrose, 5.55 mM; HEPES, 5 mM) for 2 hours at 37 °C. The EBs were pipetted gently to dissociate the cells, centrifuged (300g, 3 min) and treated with 0.25% trypsin/EDTA for 4min to obtain complete dissociation to single-cell suspensions. Cells were washed and resuspended in staining solution (PBS with 0.1% BSA) and then filtered. Antibodies were diluted in staining solution and cells were incubated on ice for 20 min. Cells were then washed and resuspended in staining solution for analysis or cell sorting.

Cell counts were collected using a LSRII (BD Biosciences) and data were analyzed using the FlowJo software. The following antibodies and dilutions were used: anti-human CD172a/b (SIRPα/β) (Biolegend; clone SE5A5; 1:200) anti-human CD13 (Biolegend; clone WM15; 1:200), anti-human CD90 (Biolegend, clone 5E10; 1:100); anti-human CD140a (PDGFRα) (Biolegend, clone 16A1; 1:50); anti-human CD309 (VEGFR2) (Biolegend, clone 7D4-6; 1:50); anti-human CD56 (NCAM) (Biolegend, clone HCD56; 1:100); anti-human CD184 (CXCR4) (Biolegend, clone 12G5; 1:100); anti-human CD117 (c-kit) (Biolegend, clone 104D2; 1:100); anti-human CD148 (Biolegend, clone A3; 1:100); HDE-1 (Keller’s lab; 1:100).

### Immunofluorescence analysis

Cells were fixed for 15 min in 4% PFA and incubated for 1 h in blocking solution (PBS with 0.1% Saponin and 1% BSA). Primary antibodies were diluted in blocking solution and incubated overnight at 4 C, followed by incubation in secondary antibody for 1 h at room temperature. Slides were then counterstained with 4,6-diamidino-2-phenylindole (DAPI) and mounted using nPG antifade mounting media. The following primary antibodies and dilutions were used: anti-cTnT (ThermoSci, clone 13-11; 1:200); anti-Mlc2v (Proteintech, 50559132; 1:200); anti-Foxa2 (Novus; NBP1-95426; 1:200); anti-Mesp1 (Aviva; ARP39374; 1:100). Secondary antibodies conjugated with Alexa dyes were obtained from Jackson Immunoresearch and used at 1:400. Fluorescence images were obtained using either Leica DM6000 or EVOS slide microscopes. Images were processed using ImageJ or Adobe Photoshop software.

### Gene expression analysis

RT-qPCR was performed as previously described (Dubois et al., 2011). Briefly, total RNA was prepared with the RNAqueous-Micro Kit (Ambion) and treated with RNase-free DNase (Ambion). Reverse transcription was performed from 100 to 500 ng of RNA using the Quanta qScript kit. Quantitative PCR was carried out on an Applied Biosystems Step One Plus using ABI SYBR Green reagents. Expression levels were normalized to the housekeeping gene TATA box binding protein (TBP). In addition, for normalization across samples genomic DNA was used to generate a standard curve. The y axis of reverse transcriptase–quantitative PCR graphs represents copy numbers of the gene of interest divided by copy numbers of TBP and therefore is an arbitrary but absolute unit that can be compared between experiments. The oligonucleotide sequences are summarized in supplementary table 2

### Statistical analysis

All experiments have a minimum of three biological replicates. qRT-PCR experiments additionally have four technical repeats for each biological replicate. Data are presented as the mean with error bars indicating the standard deviation. Statistical significance of differences between experimental groups was assessed with Student’s t-test. Differences in means were considered significant if p<0.05.

---

**Supplementary Figure 1.**
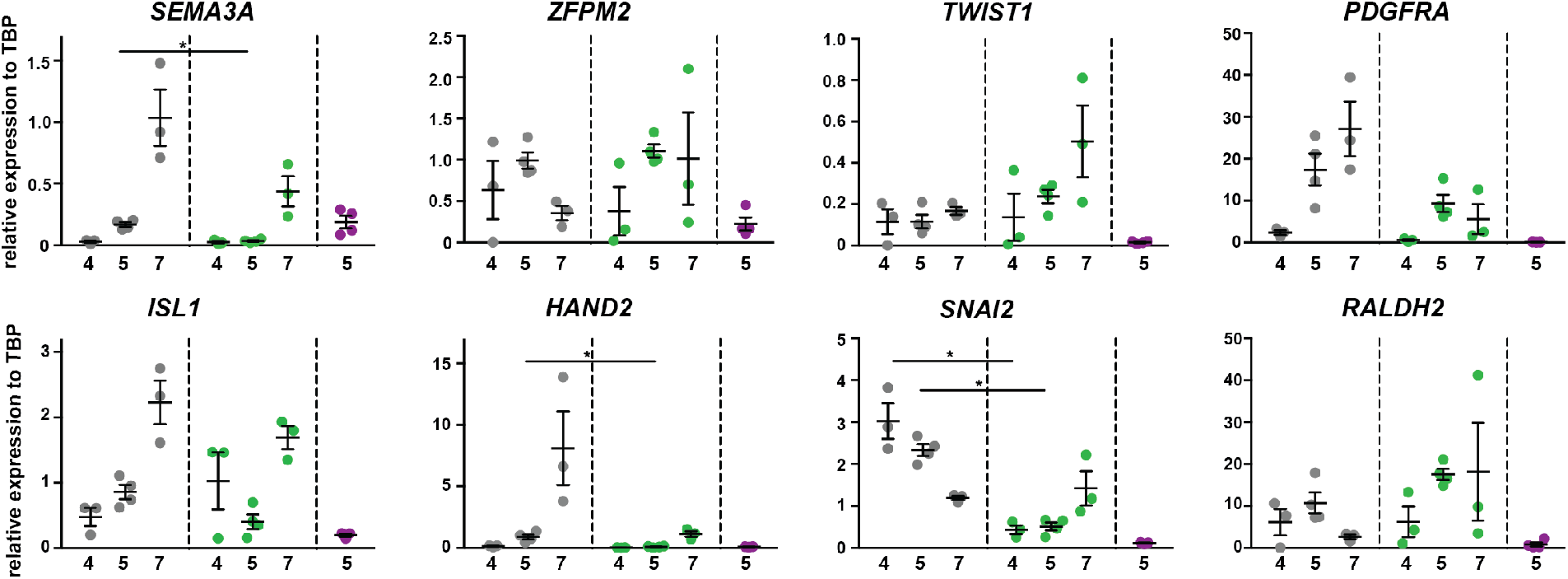
Gene expression analysis from GFP+ and GFP- cardiac mesoderm. RT-qPCR of sorted GFP- (grey) and GFP+ (green) mesoderm populations on day 4, 5 and 7 and endoderm progenitors (violet) on day 5 of previously reported mesoderm, EMT and cardiac progenitor markers. Graphs represent the relative mean expression value and SEM normalized to TBP from three independent differentiations. Statistical analysis was performed only between mesoderm populations on the same day.

**Supplementary Figure 2.**
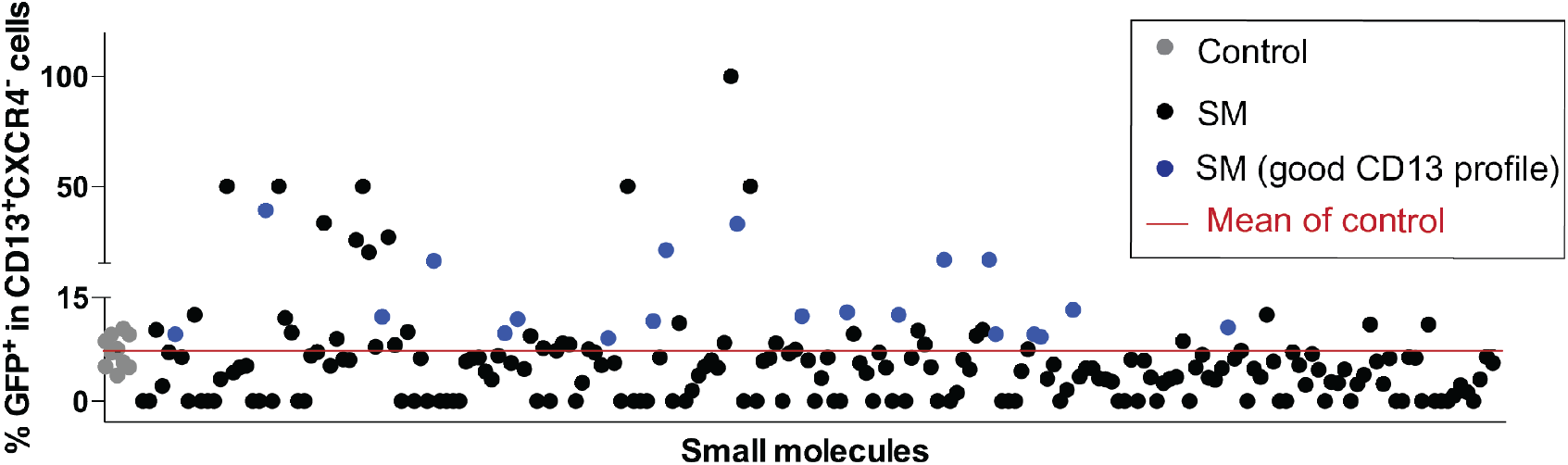
Small molecule screening to identify the pathways involved in the generation of FOXA2 mesoderm. Graphic representation of screening data from a total of 208 small molecules. Dots represent the percentages of GFP expressing cells on control (grey) or treated (black and blue) D4 mesoderm populations (CD13+CXCR4-). Small molecules that yielded similar or better mesoderm profiles on D4 when compared to control are shown in blue.

**Supplementary Figure 3.**
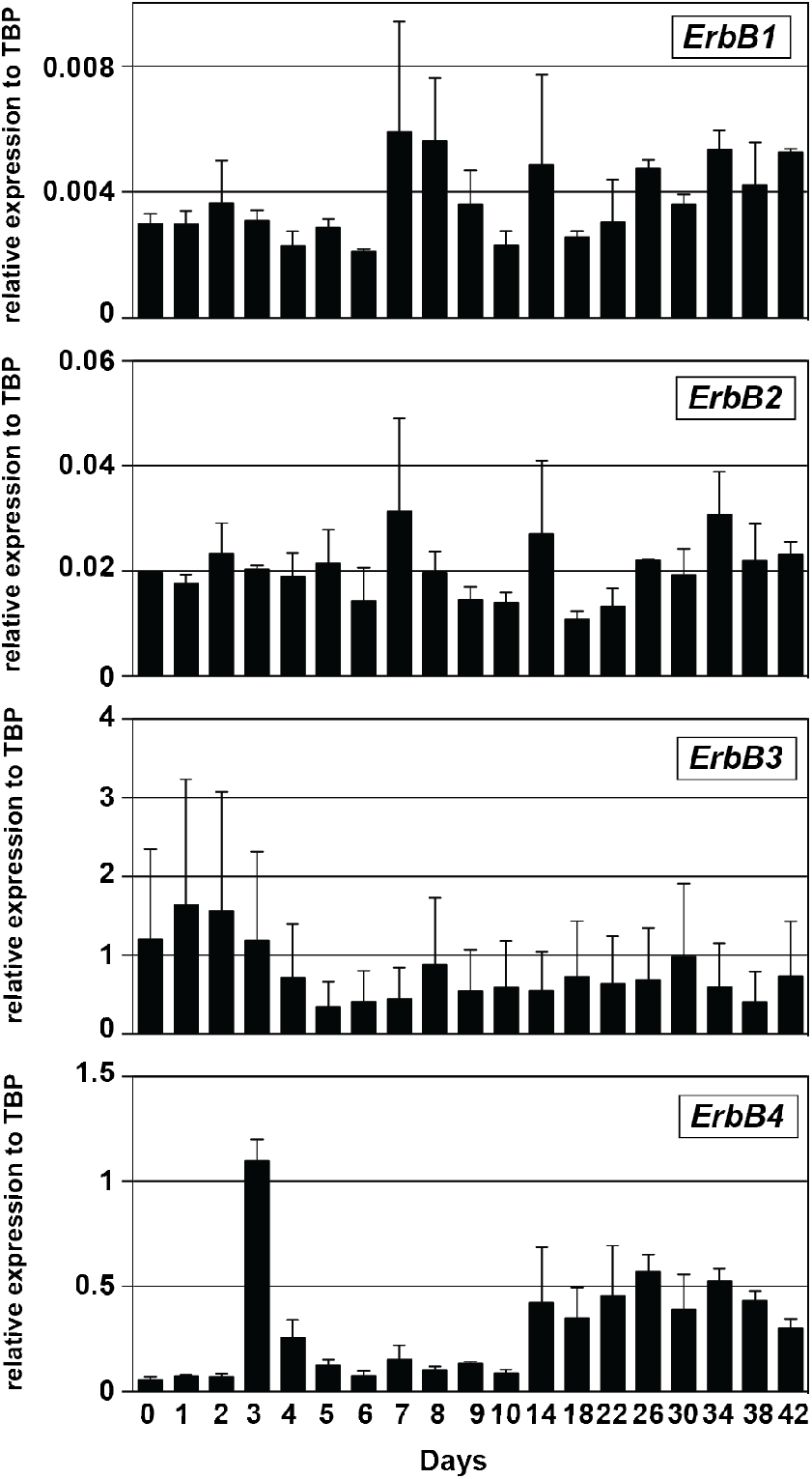
ErbB receptor expression during cardiac differentiation. ErbB3 and 4 showed increased expression during the early mesoderm differentiation, while ErbB1 and 2 showed show low and stable expression during all the differentiation.

## REFERENCES

1. Murry, C. E. & Keller, G. Differentiation of Embryonic Stem Cells to Clinically Relevant Populations: Lessons from Embryonic Development. Cell 132, 661–680 (2008).

2. Calderon, D., Bardot, E. & Dubois, N. Probing early heart development to instruct stem cell differentiation strategies. Developmental Dynamics (2016) doi:10.1002/dvdy.24441.

3. Burridge, P. W., Keller, G., Gold, J. D. & Wu, J. C. Production of de novo cardiomyocytes: Human pluripotent stem cell differentiation and direct reprogramming. Cell Stem Cell (2012) doi:10.1016/j.stem.2011.12.013.

4. Devalla, H. D., Schwach, V., Ford, J. W., Milnes, J. T., El-Haou, S., Jackson, C., Gkatzis, K., Elliott, D. A., Chuva de Sousa Lopes, S. M., Mummery, C. L., Verkerk, A. O. & Passier, R. Atrial-like cardiomyocytes from human pluripotent stem cells are a robust preclinical model for assessing atrial-selective pharmacology. EMBO Mol. Med. 7, 394–410 (2015).

5. Orlova, V. V., Van Den Hil, F. E., Petrus-Reurer, S., Drabsch, Y., Ten Dijke, P. & Mummery, C. L. Generation, expansion and functional analysis of endothelial cells and pericytes derived from human pluripotent stem cells. Nat. Protoc. (2014) doi:10.1038/nprot.2014.102.

6. Kattman, S. J., Witty, A. D., Gagliardi, M., Dubois, N. C., Niapour, M., Hotta, A., Ellis, J. & Keller, G. Stage-specific optimization of activin/nodal and BMP signaling promotes cardiac differentiation of mouse and human pluripotent stem cell lines. Cell Stem Cell 8, 228–240 (2011).

7. Dubois, N. C., Craft, A. M., Sharma, P., Elliott, D. a, Stanley, E. G., Elefanty, A. G., Gramolini, A. & Keller, G. SIRPA is a specific cell-surface marker for isolating cardiomyocytes derived from human pluripotent stem cells. Nat. Biotechnol. 29, 1011–1018 (2011).

8. Lian, X., Hsiao, C., Wilson, G., Zhu, K., Hazeltine, L. B., Azarin, S. M., Raval, K. K., Zhang, J., Kamp, T. J. & Palecek, S. P. Robust cardiomyocyte differentiation from human pluripotent stem cells via temporal modulation of canonical Wnt signaling. Proc. Natl. Acad. Sci. U. S. A. (2012) doi:10.1073/pnas.1200250109.

9. Protze, S. I., Liu, J., Nussinovitch, U., Ohana, L., Backx, P. H., Gepstein, L. & Keller, G. M. Sinoatrial node cardiomyocytes derived from human pluripotent cells function as a biological pacemaker. Nat. Biotechnol. (2017) doi:10.1038/nbt.3745.

10. Zhang, Q., Jiang, J., Han, P., Yuan, Q., Zhang, J., Zhang, X., Xu, Y., Cao, H., Meng, Q., Chen, L., Tian, T., Wang, X., Li, P., Hescheler, J., Ji, G. & Ma, Y. Direct differentiation of atrial and ventricular myocytes from human embryonic stem cells by alternating retinoid signals. Cell Res. 2011 214 21, 579–587 (2010).

11. JH, L., SI, P., Z, L., PH, B. & GM, K. Human Pluripotent Stem Cell-Derived Atrial and Ventricular Cardiomyocytes Develop from Distinct Mesoderm Populations. Cell Stem Cell 21, 179–194.e4 (2017).

12. Yang, L., Soonpaa, M. H., Adler, E. D., Roepke, T. K., Kattman, S. J., Kennedy, M., Henckaerts, E., Bonham, K., Abbott, G. W., Linden, R. M., Field, L. J. & Keller, G. M. Human cardiovascular progenitor cells develop from a KDR+ embryonic-stem-cell-derived population. Nature (2008) doi:10.1038/nature06894.

13. Vincent, S. D. & Buckingham, M. E. How to make a heart. The origin and regulation of cardiac progenitor cells. Curr. Top. Dev. Biol. 90, 1–41 (2010).

14. Meilhac, S. M., Lescroart, F., Blanpain, C. D. & Buckingham, M. E. Cardiac cell lineages that form the heart. Cold Spring Harb. Perspect. Med. 4, 13888 (2014).

15. Bruneau, B. G. The developmental genetics of congenital heart disease. Nature (2008) doi:10.1038/nature06801.

16. Devine, W. P., Wythe, J. D., George, M., Koshiba-Takeuchi, K. & Bruneau, B. G. Early patterning and specification of cardiac progenitors in gastrulating mesoderm. Elife (2014) doi:10.7554/eLife.03848.

17. Lescroart, F., Wang, X., Lin, X., Swedlund, B., Gargouri, S., Sànchez-Dànes, A., Moignard, V., Dubois, C., Paulissen, C., Kinston, S., Göttgens, B. & Blanpain, C. Defining the earliest step of cardiovascular lineage segregation by single-cell RNA-seq. Science (80-.). (2018) doi:10.1126/science.aao4174.

18. Cai, C. L., Liang, X., Shi, Y., Chu, P. H., Pfaff, S. L., Chen, J. & Evans, S. Isl1 identifies a cardiac progenitor population that proliferates prior to differentiation and contributes a majority of cells to the heart. Dev. Cell (2003) doi:10.1016/S1534-5807(03)00363-0.

19. B, S. & N, D. Of form and function: Early cardiac morphogenesis across classical and emerging model systems. Semin. Cell Dev. Biol. (2021) doi:10.1016/J.SEMCDB.2021.04.025.

20. Kinder, S. J., Loebel, D. A. F. & Tam, P. P. L. Allocation and early differentiation of cardiovascular progenitors in the mouse embryo. Trends Cardiovasc. Med. 11, 177–184 (2001).

21. Tam, P. P. L., Gad, J. M., Kinder, S. J., Tsang, T. E. & Behringer, R. R. Morphogenetic tissue movement and the establishment of body plan during development from blastocyst to gastrula in the mouse. BioEssays 23, 508–517 (2001).

22. Lescroart, F., Chabab, S., Lin, X., Rulands, S., Paulissen, C., Rodolosse, A., Auer, H., Achouri, Y., Dubois, C., Bondue, A., Simons, B. D. & Blanpain, C. Early lineage restriction in temporally distinct populations of Mesp1 progenitors during mammalian heart development. Nat. Cell Biol. 16, 829–840 (2014).

23. Chabab, S., Lescroart, F., Rulands, S., Mathiah, N., Simons, B. D. & Blanpain, C. Uncovering the Number and Clonal Dynamics of Mesp1 Progenitors during Heart Morphogenesis. Cell Rep. 14, 1–10 (2016).

24. Bardot, E., Calderon, D., Santoriello, F., Han, S., Cheung, K., Jadhav, B., Burtscher, I., Artap, S., Jain, R., Epstein, J., Lickert, H., Gouon-Evans, V., Sharp, A. J. & Dubois, N. C. Foxa2 identifies a cardiac progenitor population with ventricular differentiation potential. Nat. Commun. (2017) doi:10.1038/ncomms14428.

25. Gonzalez, D. M., Schrode, N., Ebrahim, T., Beaumont, K. G., Sebra, R. & Dubois, N. Understanding Mechanisms of Chamber-Specific Differentiation Through Combination of Lineage Tracing and Single Cell Transcriptomics. bioRxiv 2021.07.15.452540 (2021) doi:10.1101/2021.07.15.452540.

26. Joung, J. K. & Sander, J. D. TALENs: a widely applicable technology for targeted genome editing. Nat. Rev. Mol. Cell Biol. 14, 49 (2013).

27. Ogawa, S., Surapisitchat, J., Virtanen, C., Ogawa, M., Niapour, M., Sugamori, K. S., Wang, S., Tamblyn, L., Guillemette, C., Hoffmann, E., Zhao, B., Strom, S., Laposa, R. R., Tyndale, R. F., Grant, D. M. & Keller, G. Three-dimensional culture and cAMP signaling promote the maturation of human pluripotent stem cell-derived hepatocytes. Dev. 140, 3285–3296 (2013).

28. Holtzinger, A., Streeter, P. R., Sarangi, F., Hillborn, S., Niapour, M., Ogawa, S. & Keller, G. New markers for tracking endoderm induction and hepatocyte differentiation from human pluripotent stem cells. Dev. 142, 4253–4265 (2015).

29. Nostro, M. C., Sarangi, F., Ogawa, S., Holtzinger, A., Corneo, B., Li, X., Micallef, S. J., Park, I. H., Basford, C., Wheeler, M. B., Daley, G. Q., Elefanty, A. G., Stanley, E. G. & Keller, G. Stage-specific signaling through TGFβ family members and WNT regulates patterning and pancreatic specification of human pluripotent stem cells. Development 138, 861–871 (2011).

30. Green, M. D., Chen, A., Nostro, M. C., D’Souza, S. L., Schaniel, C., Lemischka, I. R., Gouon-Evans, V., Keller, G. & Snoeck, H. W. Generation of anterior foregut endoderm from human embryonic and induced pluripotent stem cells. Nat. Biotechnol. 29, 267–273 (2011).

31. Skelton, R. J. P., Brady, B., Khoja, S., Sahoo, D., Engel, J., Arasaratnam, D., Saleh, K. K., Abilez, O. J., Zhao, P., Stanley, E. G., Elefanty, A. G., Kwon, M., Elliott, D. A. & Ardehali, R. CD13 and ROR2 Permit Isolation of Highly Enriched Cardiac Mesoderm from Differentiating Human Embryonic Stem Cells. Stem Cell Reports 6, 95–108 (2016).

32. Hochgreb, T., Linhares, V. L., Menezes, D. C., Sampaio, A. C., Yan, C. Y. I., Cardoso, W. V., Rosenthal, N. & Xavier-Neto, J. A caudorostral wave of RALDH2 conveys anteroposterior information to the cardiac field. Development (2003) doi:10.1242/dev.00750.

33. Xavier-Neto, J., Rosenthal, N., Silva, F. A., Matos, T. G. F., Hochgreb, T. & Linhares, V. L. F. Retinoid signaling and cardiac anteroposterior segmentation. Genesis (2001) doi:10.1002/gene.10009.

34. Osmond, M. K., Butler, A. J., Voon, F. C. & Bellairs, R. The effects of retinoic acid on heart formation in the early chick embryo. Development 113, 1405–1417 (1991).

35. Lee, J. H., Protze, S. I., Laksman, Z., Backx, P. H., Keller, G. M., Lee, J. H., Protze, S. I., Laksman, Z., Backx, P. H. & Keller, G. M. Human Pluripotent Stem Cell-Derived Atrial and Ventricular Cardiomyocytes Develop from Distinct Mesoderm Populations Article Human Pluripotent Stem Cell-Derived Atrial and Ventricular Cardiomyocytes Develop from Distinct Mesoderm Populations. 179–194 (2017) doi:10.1016/j.stem.2017.07.003.

36. Zhang, Q., Jiang, J., Han, P., Yuan, Q., Zhang, J., Zhang, X., Xu, Y., Cao, H., Meng, Q., Chen, L., Tian, T., Wang, X., Li, P., Hescheler, J., Ji, G. & Ma, Y. Direct differentiation of atrial and ventricular myocytes from human embryonic stem cells by alternating retinoid signals. Cell Res. (2011) doi:10.1038/cr.2010.163.

37. Perl, E. & Waxman, J. S. Reiterative Mechanisms of Retinoic Acid Signaling during Vertebrate Heart Development. J. Dev. Biol. 2019, Vol. 7, Page 11 7, 11 (2019).

38. Rawnsley, D. R., Xiao, J., Lee, J. S., Liu, X., Mericko-Ishizuka, P., Kumar, V., He, J., Basu, A., Lu, M. M., Lynn, F. C., Pack, M., Gasa, R. & Kahn, M. L. The transcription factor atonal homolog 8 regulates Gata4 and friend of Gata-2 during vertebrate development. J. Biol. Chem. 288, 24429–24440 (2013).

39. Zhang, W., Shen, L., Deng, Z., Ding, Y., Mo, X., Xu, Z., Gao, Q. & Yi, L. Novel missense variants of ZFPM2/FOG2 identified in conotruncal heart defect patients do not impair interaction with GATA4. PLoS One 9, (2014).

40. Jurukovski, V. & Simon, M. Epidermal growth factor signaling pathway influences retinoid metabolism by reduction of retinyl ester hydrolase activities in normal and malignant keratinocytes. J. Cell. Physiol. 183, 265–272 (2000).

41. Takahashi, T., Takahashi, K., St. John, P. L., Fleming, P. A., Tomemori, T., Watanabe, T., Abrahamson, D. R., Drake, C. J., Shirasawa, T. & Daniel, T. O. A Mutant Receptor Tyrosine Phosphatase, CD148, Causes Defects in Vascular Development. Mol. Cell. Biol. 23, 1817–1831 (2003).

